# Association between urinary biomarkers of total sugars and sucrose intake and BMI in a cross-sectional study

**DOI:** 10.1101/099556

**Authors:** Rachel Campbell, Natasha Tasevska, Kim G Jackson, Virag Sagi-Kiss, Nick di Paolo, Jennifer S Mindell, Susan J. Lister, Kay-Tee Khaw, Gunter G. C. Kuhnle

## Abstract

Obesity is an important modifiable risk factors for chronic diseases. While there is increasing focus on the role of dietary sugars, there remains a paucity of data establishing the association between sugar intake and obesity in the general public. The objective of this study was to investigate associations of estimated sugar intake with odds for obesity in a representative samples of English adults. We used data from 434 participants of the 2005 Health Survey of England. Biomarkers for total sugar intake were measured in 24h urine samples and used to estimate intake. Linear and logistic regression analyses were used to investigate associations between estimated intake and measures of obesity (BMI, waist circumference and waist-to-hip ratio) and obesity risk., respectively. Estimated sugars intake was significantly associated with BMI, waist circumference and waist-to-hip ratio, and these associations remained significant after adjustment for estimated protein intake. Estimated sugars intake was also associated with increased odds for obesity based on BMI (OR 1.02; 95% CI 1.00; 1.04 per 10 g), waist-circumference (OR 1.03; 95% CI 1.01; 1.05) and waist-to-hip ratio (OR 1.04; 95% CI 1.02; 1.06); all OR estimates remained significant after adjusting for estimated protein intake. Our results show a significant association between biomarker-estimated total sugars intake and both measures of obesity and obesity risk, confirming positive associations between total sugar intake, measures of obesity and obesity risk. This biomarker could be used to monitor the efficacy of public health interventions.

## 1 Introduction

Dietary sugars, in particular free sugars (according to the WHO definition monosaccharides and disaccharides added to foods and beverages by the manufacturer, cook or consumer, and sugars naturally present in honey, syrups, fruit juices and fruit juice concentrates” (1)) have received increasing attention from the WHO^1^ as well as the UK government^2^ and the UK’s Scientific Advisory Committee on Nutrition (SACN)^3^. While sugar intake is often associated with an increased risk of obesity^4^, the evidence available from observational studies is more ambiguous and shows significant associations for sugar-sweetened beverages (SSB)^5,6^ only, but fails to show consistent associations for intake of sugars as nutrients ^6-9^. However, in most observational studies, sugar intake was assessed using self-report. It is likely that this has introduced bias, especially as underreporting of diet has been found to be more prevalent among obese people ^10-12^ and it is sugar-rich foods that are most commonly underreported^13^. It is possible that reporting bias contributes to the observed inverse associations between sugar intake and BMI.

Urinary sugars have been investigated^14,15^ and validated^16,17^ as dietary biomarkers of total sugars (i.e., the sum of intrinsic, milk and free sugars) and sucrose^18^ and can help to resolve the discrepancy between self-reported and true intake. This biomarker relies on the total excretion of sucrose and fructose within 24h and therefore requires complete 24h urine samples. While we have been able to show a positive association between the biomarker measured in spot urines and BMI^16,19^, the lack of validation data on the performance of sucrose and fructose as dietary biomarkers from spot urines weakens these results.

In this study, we have investigated the association between sugar intake and obesity risk using exclusively nutritional biomarkers and not relying on self-reported data. The results of this study will allow us to test the feasibility of applying this biomarker to an existing cohort as an instrument to monitor consumption and to investigate associations between sugar intake and obesity.

## 2 Method

### 2.1 Study population

The Health Survey for England is a health examination survey of nationally-representative samples of the general population. A new, random, household-based sample has been selected annually since 1991. Individuals living at the selected private addresses are recruited to the study, answer a questionnaire through face-to-face interview, and have trained interviewers measure height and weight. Nurses take other physical measurements and collect biological samples ^20^. The measurement of height, weight (interviewer), and waist and hip circumference (nurse) followed the protocols of the 2003 Health Survey for England ^21^. No data on diet, except for fruit and vegetable intake, were collected by interview.

We used data of participants from the 2005 Health Survey for England (HSE 2005) with the aim of obtaining a nationally-representative sample of the general population aged 19 to 64 years living in England. As a supplement to the main HSE 2005, a sub-sample of adult participants were asked to provide a 24-hour urine sample. Overall, 498 survey participants (200 men, 298 women, Table 1), aged 19 and over, who provided a 24-hour urine sample were identified and included in the study. Data collection took place between October 2005 and July 2006, with the majority of fieldwork being completed by March 2006. If more than one 24-h urine sample was available for one participant, the first sample was used.

**Table 1:**
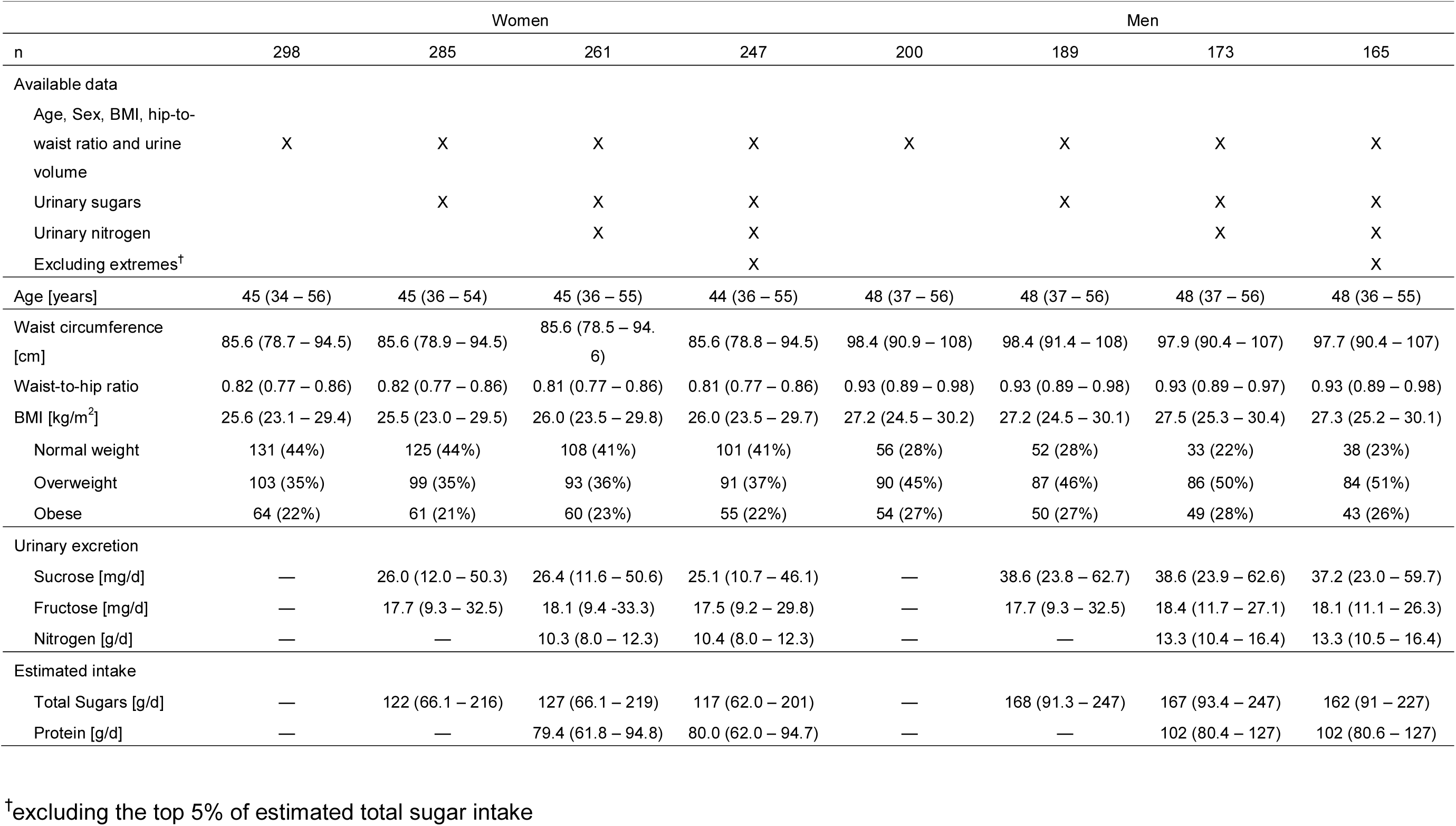
Study population characteristics and description of analytical sample. Median and inter-quartile range or absolute number and proportion.

|  | Women |  |  |  | Men |  |  |  |
| --- | --- | --- | --- | --- | --- | --- | --- | --- |
| n | 298 | 285 | 261 | 247 | 200 | 189 | 173 | 165 |
| Available data |  |  |  |  |  |  |  |  |
| Age, Sex, BMI, hip-to-waist ratio and urine volume | X | X | X | X | X | X | X | X |
| Urinary sugars |  | X | X | X |  | X | X | X |
| Urinary nitrogen |  |  | X | X |  |  | X | X |
| Excluding extremes<sup>†</sup> |  |  |  | X |  |  |  | X |
| Age [years] | 45 (34 – 56) | 45 (36 – 54) | 45 (36 – 55) | 44 (36 – 55) | 48 (37 – 56) | 48 (37 – 56) | 48 (37 – 56) | 48 (36 – 55) |
| Waist circumference [cm] | 85.6 (78.7 – 94.5) | 85.6 (78.9 – 94.5) | 85.6 (78.5 – 94.6) | 85.6 (78.8 – 94.5) | 98.4 (90.9 – 108) | 98.4 (91.4 – 108) | 97.9 (90.4 – 107) | 97.7 (90.4 – 107) |
| Waist-to-hip ratio | 0.82 (0.77 – 0.86) | 0.82 (0.77 – 0.86) | 0.81 (0.77 – 0.86) | 0.81 (0.77 – 0.86) | 0.93 (0.89 – 0.98) | 0.93 (0.89 – 0.98) | 0.93 (0.89 – 0.97) | 0.93 (0.89 – 0.98) |
| BMI [kg/m<sup>2</sup>] | 25.6 (23.1 – 29.4) | 25.5 (23.0 – 29.5) | 26.0 (23.5 – 29.8) | 26.0 (23.5 – 29.7) | 27.2 (24.5 – 30.2) | 27.2 (24.5 – 30.1) | 27.5 (25.3 – 30.4) | 27.3 (25.2 – 30.1) |
| Normal weight | 131 (44%) | 125 (44%) | 108 (41%) | 101 (41%) | 56 (28%) | 52 (28%) | 33 (22%) | 38 (23%) |
| Overweight | 103 (35%) | 99 (35%) | 93 (36%) | 91 (37%) | 90 (45%) | 87 (46%) | 86 (50%) | 84 (51%) |
| Obese | 64 (22%) | 61 (21%) | 60 (23%) | 55 (22%) | 54 (27%) | 50 (27%) | 49 (28%) | 43 (26%) |
| Urinary excretion |  |  |  |  |  |  |  |  |
| Sucrose [mg/d] | — | 26.0 (12.0 – 50.3) | 26.4 (11.6 – 50.6) | 25.1 (10.7 – 46.1) | — | 38.6 (23.8 – 62.7) | 38.6 (23.9 – 62.6) | 37.2 (23.0 – 59.7) |
| Fructose [mg/d] | — | 17.7 (9.3 – 32.5) | 18.1 (9.4 -33.3) | 17.5 (9.2 – 29.8) | — | 17.7 (9.3 – 32.5) | 18.4 (11.7 – 27.1) | 18.1 (11.1 – 26.3) |
| Nitrogen [g/d] | — | — | 10.3 (8.0 – 12.3) | 10.4 (8.0 – 12.3) | — | — | 13.3 (10.4 – 16.4) | 13.3 (10.5 – 16.4) |
| Estimated intake |  |  |  |  |  |  |  |  |
| Total Sugars [g/d] | — | 122 (66.1 – 216) | 127 (66.1 – 219) | 117 (62.0 – 201) | — | 168 (91.3 – 247) | 167 (93.4 – 247) | 162 (91 – 227) |
| Protein [g/d] | — | — | 79.4 (61.8 – 94.8) | 80.0 (62.0 – 94.7) | — | — | 102 (80.4 – 127) | 102 (80.6 – 127) |

<sup>†</sup>excluding the top 5% of estimated total sugar intake

### 2.2 24-hour urine collection

Participants were asked to collect all urine they passed during a 24-hour period starting from the second morning urine void of the 24-hour collection day, and ending with the first urine void the following morning. P-amino-benzoic acid (PABA) was used to test for completeness of 24h urine collection and only complete samples (with >85% PABA recovery in urine) were used for this analysis^22^. All samples were stored at -20°C until analysis.

### 2.3 Analysis of urinary sucrose and fructose

Urine samples were thawed at room temperature, centrifuged to remove protein aggregates and analysed using an ILAB600 clinical chemistry analyser (Werfen (UK) Limited, Warrington) with a sucrose, fructose and glucose enzyme kit (Sucrose/D-Glucose/D-Fructose; Boehringer Mannheim, R-Biopharm, Enzymatic BioAnalysis/Food Analysis, Darmstadt, Germany). This method determines D-glucose by measuring NADPH + H^+^ formation following phosphorylation of D-glucose by hexokinase and subsequent oxidation by NADPH+-dependent glucose-6-phosphate dehydrogenase. NADPH + H^+^ is determined by changes in absorption at 340 nm. Sucrose and D-fructose are determined indirectly following the conversion of D-fructose into D-glucose by phosphoglucose-isomerase or β-fructosidase and calculating the difference in D-glucose concentration before and after conversion. The concentration range for sucrose and D-fructose was 2.5 to 200 mg/L, for D-glucose it was 2.5 to 150 mg/L; samples exceeding these concentrations were diluted 1 in 10 with purified water and reanalysed. The intra-assay CV for a 25 mg/L glucose quality control (QC) sample was less than 2% and the inter-assay CV was 3.6%. The inter-assay CV was also determined for fructose and sucrose and found to be less than 7%. All concentrations measured were above the lower-limit of quantification. 24-h urinary sucrose and fructose were calculated based on urinary fructose and sucrose concentration (mg/L) and 24-h urine volume.

### 2.4 Analysis of urinary nitrogen

We measured 24-h urinary nitrogen, a recovery biomarker for protein intake, to partialy control for non-sugars energy intake. Urine samples were thawed at room temperature prior to analysis. Approximately 1 ml of samples was weighed into a tin foil capsule. For Total Nitrogen (N %) determination, the sample was combusted in oxygen and the nitrogen released measured with a thermal conductivity cell using a LECO FP-428 Analyser (LECO Corp., St. Joseph, MI). The coefficient of variation for within-run and within-laboratory precision was 1.77 and 3.80 %, respectively for an internal quality control sample containing 1 % N. The limit of quantification for the test was 0.018 % N.

### 2.5 Biomarker-based estimates of total sugars and protein intake

Estimated total sugars intake was calculated based on a calibration equation for the sugars biomarker developed from a feeding study conducted in the UK (16), which describes the association between the biomarker and true intake^17^

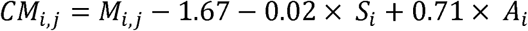

where CM is log transformed calibrated biomarker of person i at time point j, i.e. predicted total sugars intake, M is log transformed sum of 24-hour urine fructose and sucrose, S is sex (male: S=0, female: S=1) and A is log transformed age. Estimated protein intake was calculated based on the assumptions that 81% of dietary nitrogen is recovered from urine^23^ and an average nitrogen content of proteins is 16% [P: protein intake (g/d), N: total nitrogen excretion (g/d)]:

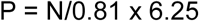

### 2.6 Data handling

Calculated fructose and sucrose concentrations of zero were assigned a value of 0.1 to allow for a log_2_-transformation of the data.

### 2.7 Statistical analyses

All data were processed using R^24^ version 3.3.2. Urinary fructose, urinary sucrose and the sum of 24-hour urinary fructose and sucrose were skewed to the right and log_2_-transformed, whereas biomarker estimates of total sugars and protein intakes were used without transformation. The ratio of urinary sugars and estimated sugar intake, to urinary nitrogen and estimated protein intake, respectively, were log_2_-transformed. We used the ratios of estimated total sugars to protein intake or urinary sugars to urinary nitrogen to investigate the effect of sugars while controlling for dietary composition. Unadjusted models were used when investigating associations between estimated total sugars intake and BMI and obesity risk (based on WHO definition either as BMI ≥ 30 kg/m^2^ or waist-to-hip ratio > 0.85 for women and > 0.90 for men), given the calibration equation for the sugars biomarker which we used to estimate total sugars intake included age and sex. Models with uncalibrated urinary fructose, uncalibrated urinary sucrose or estimated protein intake and BMI and obesity risk were adjusted for age and sex. Associations with BMI were investigated using linear regression models; OR for obesity (as estimate of risk) was estimated using logistic regression. Urinary nitrogen or estimated protein intake was included in the models to control for protein intake as a contributor to energy intake. P<0.05 was used as threshold for statistical significance.

## 3 Results

### 3.1 Study population

Study population characteristics and description of the analytical sample are shown in Table 1. Complete data on age, sex, BMI, waist-to-hip-ratio and 24h urine volume were available for 298 women and 200 men (n=498). Due to missing samples or insufficient volume, not all samples could be analysed for urinary biomarkers; data on urinary sugars and nitrogen are available for 261 women and 173 men only (n=434).

The distribution of estimated dietary sugar intake (median 144 g/d, range 0 – 2777 g/d) was skewed right with some extremely high values. We have therefore truncated the data at the 95^th^ centile of estimated intake (527 g/d). The remaining sample included 247 women and 165 men (n=411). Participants in the top 5^th^ centile (14 women, 8 men) were older (mean age 50.6 years *vs* 44.8 years, *t*-test: p=0.024) and had a higher excretion of sucrose (247 mg/d *vs* 36.4 mg/d, *t*-test: p<0.001) and fructose (84.9 mg/d *vs* 22.3 mg.d, *t*-test: p<0.001) than those in the remaining sample. There were however no statistically significant differences in BMI, waist circumference, waist-to-hip ratio or protein intake.

### 3.2 Associations between biomarker excretion, measures of obesity and obesity risk

Associations between urinary fructose, urinary sucrose, the sum of 24-hour urinary fructose and sucrose, and 24-hour urinary nitrogen and measures of obesity (BMI, waist circumference and waist-to-hip ratio), adjusted for age and sex, are shown in Table 2. We found a significant positive association for 24h urinary sucrose with all measures of obesity; these associations were strengthened when including 24h urinary fructose and 24h urinary nitrogen in the model. Total urinary sugars were significantly associated only with waist circumference and waist-to-hip ratio, although the former association became non-significant after adjusting for urinary nitrogen. There were no associations between any marker and total urinary fructose, whereas total urinary nitrogen was significantly associated with BMI and waist circumference, but not waist-to-hip ratio.

**Table 2:** Associations between 24h excretion of sucrose, fructose and nitrogen and BMI, waist-circumference and waist-to-hip-ratio (β and 95% CI). Data were log2-transformed and models are adjusted for age and sex. Estimates in each column represent a separate model.

| Regression coefficient ($\beta$ and 95% CI per doubling of excretion) | | | | | | | | | |
| --- | --- | --- | --- | --- | --- | --- | --- | --- | --- |
| BMI [kg/m<sup>2</sup>] |  |  |  |  |  |  |  |  |  |
| Sum of 24-h urinary sucrose and fructose [mg/d] | 0.201<br>(-0.099; 0.501) | — | — | — | — | 0.171<br>(-0.129; 0.471) | — | — | — |
| 24-h urinary sucrose [mg/d] | — | 0.243<br>(0.029; 0.458)<sup>†</sup> | — | 0.309<br>(0.073; 0.545)<sup>†</sup> | — | — | 0.236<br>(0.023; 0.450)<sup>†</sup> | — | 0.318<br>(0.083; 0.553)<sup>††</sup> |
| 24-h urinary fructose [mg/d] | — | — | -0.030<br>(-0.258; 0.199) | -0.166<br>(-0.416; 0.083) | — | — | — | -0.067<br>(-0.296; 0.163) | -0.209<br>(-0.460; 0.042) |
| 24-h urinary Nitrogen [g/d] | — | — | — | — | 0.960<br>(0.077; 1.842)<sup>†</sup> | 0.910<br>(0.024; 1.797)<sup>†</sup> | 0.931<br>(0.052; 1.810)<sup>†</sup> | 0.997<br>(0.105; 1.890)<sup>†</sup> | 1.038<br>(0.151; 1.925)<sup>†</sup> |
| Waist circumference [cm] |  |  |  |  |  |  |  |  |  |
| Sum of 24-h urinary sucrose and fructose [mg/d] | 0.754<br>(0.022; 1.486)<sup>†</sup> | — | — | — | — | 0.666<br>(-0.066; 1.397) | — | — | — |
| 24-h urinary sucrose [mg/d] | — | 0.782<br>(0.259; 1.305)<sup>†</sup><br>† | — | 0.946<br>(0.370; 1.521)<sup>††</sup> | — | — | 0.762<br>(0.242; 1.282)<sup>††</sup> | — | 0.973<br>(0.402; 1.544)<sup>†††</sup> |
| 24-h urinary fructose [mg/d] | — | — | 0.004<br>(-0.556; 0.563) | -0.415<br>(-1.024; 0.194) | — | — | — | -0.105<br>(-0.666; 0.456) | -0.540<br>(-1.150; 0.070) |
| 24-h urinary Nitrogen [g/d] | — | — | — | — | 2.857<br>(0.702; 5.012)<sup>†</sup><br>† | 2.665<br>(0.506; 4.825)<sup>†</sup> | 2.764<br>(0.626; 4.901)<sup>††</sup> | 2.916<br>(0.736; 5.096)<sup>††</sup> | 3.041<br>(0.887; 5.196)<sup>††</sup> |
| Waist-to-hip ratio [x 100] |  |  |  |  |  |  |  |  |  |
| Sum of 24-h urinary sucrose and fructose [mg/d] | 0.474<br>(0.081; 0.867)<sup>†</sup> | — | — | — | — | 0.458<br>(0.063; 0.853)<sup>††</sup> | — | — | — |
| 24-h urinary sucrose [mg/d] | — | 0.415<br>(0.133; 0.696)<sup>††</sup> | — | 0.496<br>(0.186; 0.805)<sup>††</sup> | — | — | 0.411<br>(0.129; 0.692)<sup>††</sup> | — | 0.502<br>(0.192; 0.812)<sup>††</sup> |
| 24-h urinary fructose<br>[mg/d] | — | — | 0.015<br>(-0.286; 0.316) | -0.204<br>(-0.532; 0.123) | — | — | — | -0.008<br>(-0.312; 0.296) | -0.233<br>(-0.564; 0.098) |
| 24-h urinary Nitrogen [g/d] | — | — | — | — | 0.627<br>(-0.540; 1.794) | 0.495<br>(-0.672; 1.662) | 0.577<br>(-0.581; 1.734) | 0.632<br>(-0.549; 1.813) | 0.697<br>(-0.472; 1.865) |

<sup>†</sup>p<0.05; <sup>††</sup>p<0.01, <sup>†††</sup>p<0.001

Total urinary sucrose was positively associated with obesity risk when using waist circumference as the obesity marker, and this association became stronger when adjusting for 24h urinary fructose, and for both 24h urinary fructose and 24h urinary nitrogen (Table 3). Total urinary sucrose was also positively associated with obesity risk when using waist-to-hip-ratio as obesity marker, but only after adjustment for urinary fructose and urinary fructose and nitrogen. We found no statistically significant increase in obesity risk when using BMI as the obesity marker.

**Table 3:** Associations between 24h uriny excretion of sucrose, fructose and nitrogen and and odds for obesity (OR and 95% CI). Data were log2-transformed and models are adjusted for age and sex. Estimates in each column represent a separate model.

| Obesity risk (OR and 95% CI – per doubling of excretion) |  |  |  |  |  |  |  |  |  |
| --- | --- | --- | --- | --- | --- | --- | --- | --- | --- |
| BMI ≥ 30 kg/m<sup>2</sup> |  |  |  |  |  |  |  |  |  |
| Sum of 24-h urinary sucrose and fructose [mg/d] | 1.04<br>(0.90; 1.22) | — | — | — | — | 1.03<br>(0.89; 1.21) | — | — | — |
| 24-h urinary sucrose [mg/d] | — | 1.08<br>(0.97; 1.22) | — | 1.12<br>(0.98; 1.27) | — | — | 1.08<br>(0.97; 1.22) | — | 1.12<br>(0.99; 1.28) |
| 24-h urinary fructose [mg/d] | — | — | 0.99<br>(0.89; 1.10) | 0.94<br>(0.84; 1.07) | — | — | — | 0.97<br>(0.87; 1.09) | 0.93<br>(0.82; 1.05) |
| 24-h urinary Nitrogen [g/d] | — | — | — | — | 1.53<br>(0.97; 2.48) | 1.52<br>(0.96; 2.46) | 1.53<br>(0.97; 2.48) | 1.56<br>(0.99; 2.53) | 1.60<br>(1.00; 2.61) |
| Waist circumference > 85 cm (women) or 94 cm (men) |  |  |  |  |  |  |  |  |  |
| Sum of 24-h urinary sucrose and fructose [mg/d] | 1.04 (0.90; 1.22) | — | — | — | — | 1.13<br>(0.99; 1.30) | — | — | — |
| 24-h urinary sucrose [mg/d] | — | 1.12<br>(1.02; 1.24)<sup>†</sup> | — | 1.16<br>(1.04; 1.29)<sup>†</sup> | — | — | 1.12<br>(1.02; 1.24)<sup>†</sup> | — | 1.16<br>(1.04; 1.30)<sup>†</sup> |
| 24-h urinary fructose [mg/d] | — | — | 0.99<br>(0.89; 1.10) | 0.93<br>(0.82; 1.04) | — | — | — | 0.97<br>(0.87; 1.09) | 0.91<br>(0.80; 1.03) |
| 24-h urinary Nitrogen [g/d] | — | — | — | — | 1.43<br>(0.97; 2.15) | 1.38<br>(0.93; 2.07) | 1.41<br>(0.95; 2.12) | 1.46<br>(0.98; 2.19) | 1.49<br>(0.99; 2.25) |
| Waist-to-hip ratio > 0.85 (women) or 0.90 (men) |  |  |  |  |  |  |  |  |  |
| Sum of 24-h urinary sucrose and fructose [mg/d] | 1.07<br>(0.93; 1.24) | — | — | — | — | 1.07<br>(0.93; 1.24) | — | — | — |
| 24-h urinary sucrose [mg/d] | — | 1.11<br>(1.00; 1.24) | — | 1.15<br>(1.03; 1.30)<sup>†</sup> | — | — | 1.11<br>(1.00; 1.24) | — | 1.16<br>(1.03; 1.31)<sup>†</sup> |
| 24-h urinary fructose [mg/d] | — | — | 0.97<br>(0.87; 1.08) | 0.91<br>(0.80; 1.03) | — | — | — | 0.97<br>(0.87; 1.08) | 0.90<br>(0.80; 1.02) |
| 24-h urinary Nitrogen [g/d] | — | — | — | — | 1.09<br>(0.71; 1.68) | 1.08<br>(0.70; 1.66) | 1.09<br>(0.71; 1.67) | 1.11<br>(0.72; 1.71) | 1.13<br>(0.74; 1.76) |

<sup>†</sup>p<0.05

### 3.3 Associations between estimated intake, measures of obesity and obesity risk

Estimated total sugars and protein intake were positively associated with BMI, waist circumference and waist-to-hip ratio, both independently and when combined in the same model (Table 4; Fig 1). They were also positively associated with obesity risk when using waist-to-hip-ratio as the obesity marker. However, associations were weaker for BMI and waist circumference. Significant associations were observed only for estimated protein intake (both independently and in the combined model) and estimated sugar intake when using BMI as the obesity marker, and only for estimated sugar intake in the combined model when using waist circumference.

**Fig 1:**
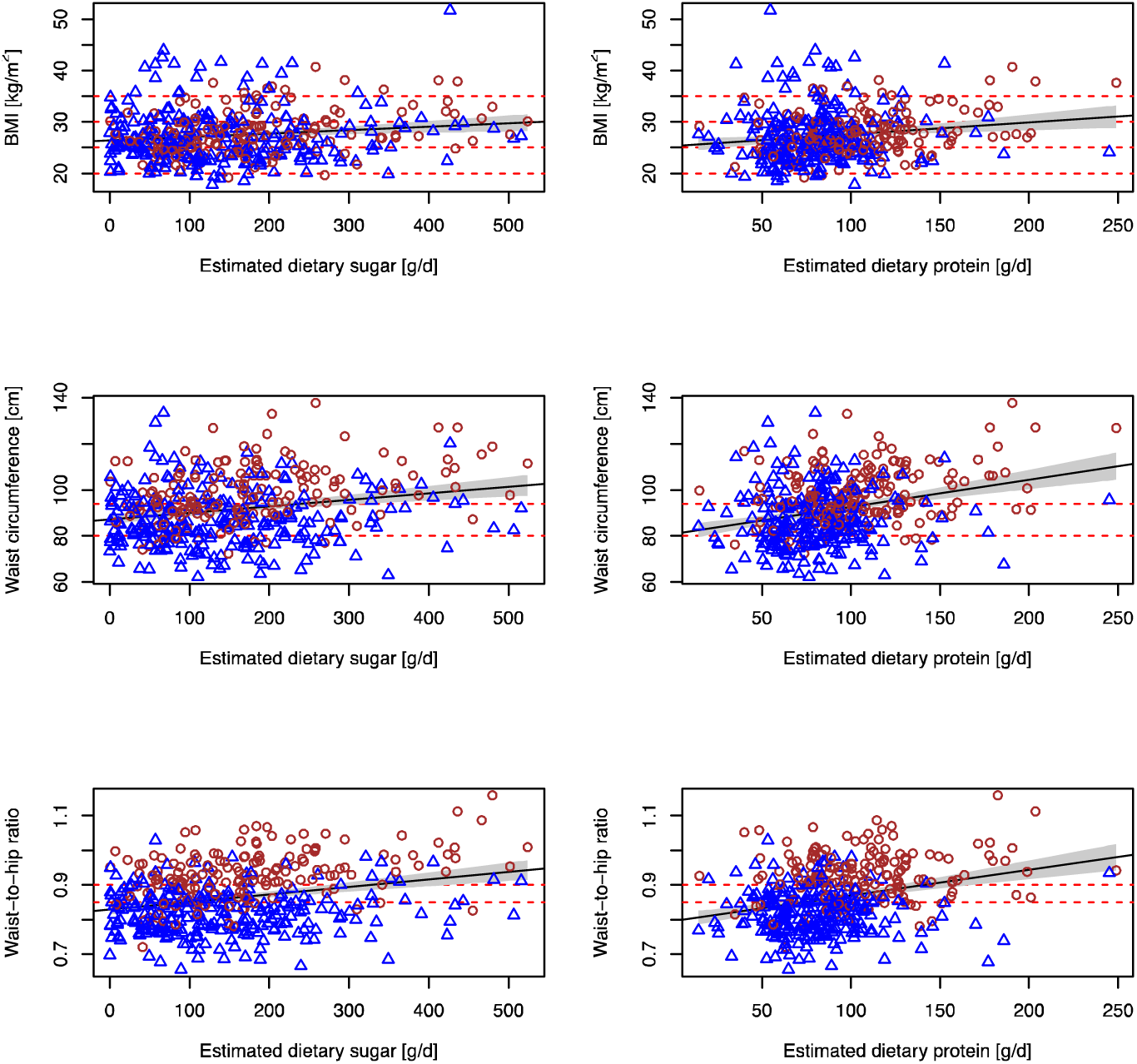
Associations between estimated sugars and protein intake and obesity markers. Associations between estimated sugars and protein intake and BMI, waist circumference and waist-to-hip ratio in men (brown circles) and women (blue triangles)

**Table 4:** Associations between estimated total sugars and protein intake and BMI (β and 95% CI per 10 g) and obesity risk (OR and 95% CI per 10 g). Estimates in each column represent a separate model.

|  | Marker of obesity |  |  |  |  |  |  |  |  |
| --- | --- | --- | --- | --- | --- | --- | --- | --- | --- |
|  | BMI |  |  | Waist circumference |  |  | Waist-to-hip ratio [x 100] |  |  |
|  | Regression coefficient (β and 95% CI per 10 g/d) |  |  |  |  |  |  |  |  |
| Total estimated sugars intake [10 g/d] | 0.066<br>(0.024; 0.108)<sup>††</sup> | — | 0.055<br>(0.012; 0.097)<sup>†</sup> | 0.281<br>(0.165; 0.396)<sup>†††</sup> | — | 0.220<br>(0.106; 0.333)<sup>†††</sup> | 0.219<br>(0.145; 0.292)<sup>†††</sup> | — | 0.182<br>(0.109; 0.254)<sup>†††</sup> |
| Estimated protein intake [10 g/d] | — | 0.229<br>(0.095; 0.365)<sup>†††</sup> | 0.197<br>(0.061; 0.333)<sup>††</sup> | — | 1.180<br>(0.818; 1.543)<sup>†††</sup> | 1.049<br>(0.686; 1.412)<sup>†††</sup> | — | 0.745<br>(0.511; 0.979)<sup>†††</sup> | 0.636<br>(0.404; 0.868)<sup>†††</sup> |
|  | Obesity risk<sup>‡</sup> (OR and 95% CI per 10 g/d) |  |  |  |  |  |  |  |  |
| Total estimated sugars intake [10 g/d] | 1.02<br>(1.00; 1.04)<sup>†</sup> | — | 1.02<br>(1.00; 1.04) | 1.03<sup>††</sup><br>(1.01; 1.05) | — | 1.03<br>(1.01; 1.05)<sup>††</sup> | 1.04<br>(1.02; 1.06)<sup>†††</sup> | — | 1.03<br>(1.01; 1.05)<sup>†††</sup> |
| Estimated protein intake [10 g/d] | — | 1.08<br>(1.02; 1.15)<sup>†</sup> | 1.07<br>(1.01; 1.14)<sup>†</sup> | — | 1.05<br>(0.99; 1.12) | 1.03<br>(0.97; 1.10) | — | 1.12<br>(1.05; 1.19)<sup>†††</sup> | 1.10<br>(1.03; 1.17)<sup>††</sup> |

<sup>†</sup>p<0.05; <sup>††</sup>p<0.01; <sup>†††</sup>p<0.001; <sup>‡</sup> BMI ≥ 30 kg/m<sup>2</sup>; waist circumference > 85 cm (women) or 94 cm (men); waist-to-hip ratio > 0.85 (women) or 0.90 (men)
(women) or 0.90 (men)

### 3.4 Associations between ratio of sugar-to-protein intake, BMI and obesity risk

We investigated the ratios of estimated total sugars to protein intake or urinary sugars to urinary nitrogen to investigate the effect of sugar intake while controlling for dietary composition (Table 5). These data showed a positive association between the urinary sucrose-to-nitrogen ratio and measures of obesity, especially after adjustment for urinary fructose-to-nitrogen ratio. The increase in urinary sucrose to nitrogen ratio was associated with statistically significant increased risk of obesity (waist circumference and waist-to-hip ratio) after adjusting for urinary fructose to nitrogen ratio in the model. We found no statistically significant increase in obesity risk with estimated total sugars to nitrogen intake, urinary sugars to nitrogen ratio or urinary fructose to nitrogen ratio.

**Table 5:**
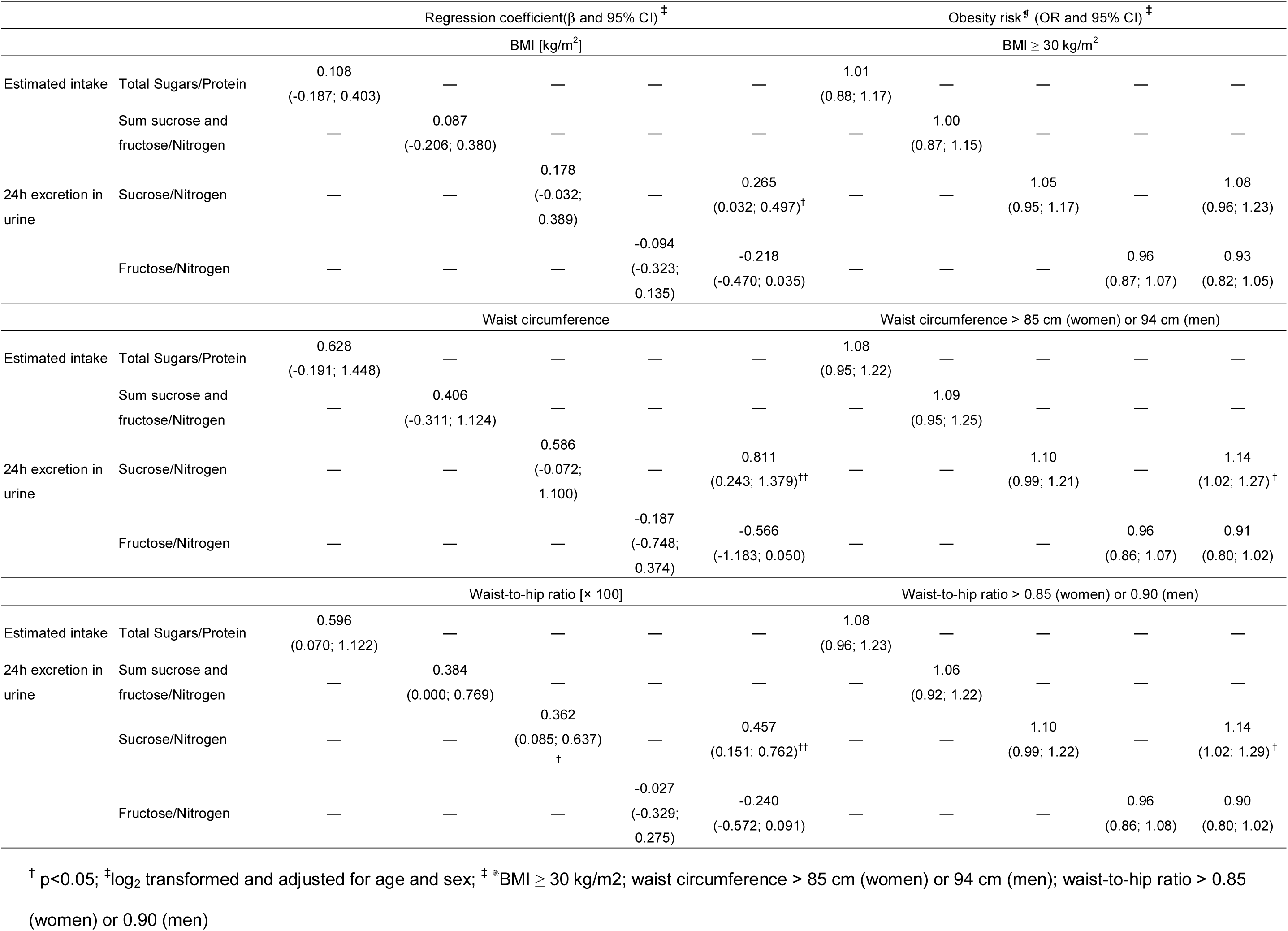
Associations between ratio of sugars and protein intake, and ratio of urinary sugars and nitrogen and BMI (β and 95% CI) and obesity risk (OR and 95% CI). Estimates in each column represent a separate model.

## 4 Discussion

In this study, we have used exclusively biomarker and biomarker-estimated data and not self-reported data to investigate associations between sugar intake and obesity risk. In our study population, using biomarker-based intake estiamtes, sugars were significantly associated with BMI, waist circumference and waist-to-hip ratio, and these associations remained significant after adjustment for biomarker-based protein intake. Estimated sugars intake was also associated with increased odds for obesity as measured by BMI, waist-circumference and waist-to-hip ratio. The association between sugar intake and obesity risk in the general public is difficult to investigate because of the known limitation of self-reported dietary assessment, in particular the tendency to underreport the intake of perceived unhealthy foods and foods with high sugar content, especially among overweight individuals^11^. Indeed, observational studies relying on self-reported intake have long produced inconsistent results and generated controversy. Consistent data are available only for an association between obesity and sweetened beverages but not total sugar intake^9^. The objective assessment of sugar intake using a dietary biomarker^16,17^ relies on total daily sucrose and fructose excretion and therefore the availability of 24h urine samples, which are often not available. Previously, this biomarker has been adapted for use with spot urine samples, showing a significant association between sucrose intake and BMI in two subsets of a cohort study, EPIC Norfolk^4,19^. However, while the biomarker measured in 24-h urine samples has been thoroughly validated, this is not the case for the biomarker measured in spot urine samples. Controlled feeding studies are needed to investigate and characterize the use of sucrose and fructose from spot urine as a biomarker for sugars^18^.

Total sugars and protein intake in our women (117 g/d, 80 g/d) and men (162 g/d; 102 g/d) estimated using biomarkers was higher than in the 2008/9 UK National Diet and Nutrition Survey (NDNS) (women: 78 g/d men, 66 g/d: men: 107 g/d, 89 g/d)^25^. An explanation for this discrepancy, in particular for total sugars intake, is that the NDNS relies on self-reported data and underreporting, in particular of sugar intake, is likely.

Our data showed a significant association between biomarker-estimated total sugar intake and both measures of obesity and obesity risk, confirming positive associations between total sugar intake, measures of obesity and obesity risk. The main strengths of this study are that the samples are a representative selection of the English population and that 24h urine samples were available. Moreover, the calibration equation that was used to calibrate the biomarker and generate estimate of sugars intake was developed in a UK feeding study under a UK diet. Even though no data on energy intake were available in the study, we were able to partially control for non-sugars energy using an objective measure of protein intake. Limitations include the small sample size; many associations were of borderline statistical significance and a larger study would allow further exploration. Furthermore, the application of biomarkers assumes an equilibrium, i.e., that participants do not change their body composition^26^, which information was not available, and could have introduced bias.

There was no information about stomach ulcers – which increase gastrointestinal permeability for (unhydrolized) sucrose – or impaired kidney function, for example as a result of type 2 diabetes, which could affect urinary fructose and sucrose excretion. Previous research has shown that neither obesity nor stomach ulcers have a significant impact on the biomarker used^19,27^, but there is a paucity of data investigating the effect of impaired renal function. As sucrose is excreted rapidly and almost completely in urine^28^, it is unlikely that diabetic kidney disease affects urinary sucrose concentrations. The physiological processes are more complex for fructose as it involves active reabsorption in the kidney^29^ and higher urinary fructose concentrations have been observed in patients with diabetes^30^, although it is not clear whether this is due to impaired kidney function. This would result in an overestimation of sugar intake in those participants.

While BMI is commonly used to diagnose obesity, there are some limitations due to its inability to discriminate between fat and lean mass^31^. We have therefore also included waist circumference and waist-to-hip ratio in our analyses and the results are comparable. Indeed, associations between estimated sugar intake and obesity risk are stronger when using waist-circumference and waist-to-hip ratio as measures of obesity.

A possible explanation for the association between sugar intake and measures of obesity could be that sugar intake is an important contributor of energy intake. The paucity of validated recovery biomarkers for fat and total carbohydrate intake makes it difficult to assess total energy intake without double labelled water^32^ or retrospectively. Protein is the only macronutrient for which intake can be estimated reliably with a dietary biomarker, total urinary nitrogen excretion^12,23^. In the UK, protein contributed approximately 15% to 20% of total daily energy intake^25^ and we have therefore used biomarker-estimated protein intake to partially adjust for non-sugar energy intake. Urinary nitrogen excretion was positively associated with BMI and waist circumference, but not waist-to-hip ratio. Independently, estimated protein intake was also associated with many measures of obesity and obesity risk based on BMI and waist-to-hip rato. These associations remained significant when combining sugar and protein in the same model, although both became slightly attenuated (Fig 2).

**Fig 2:**
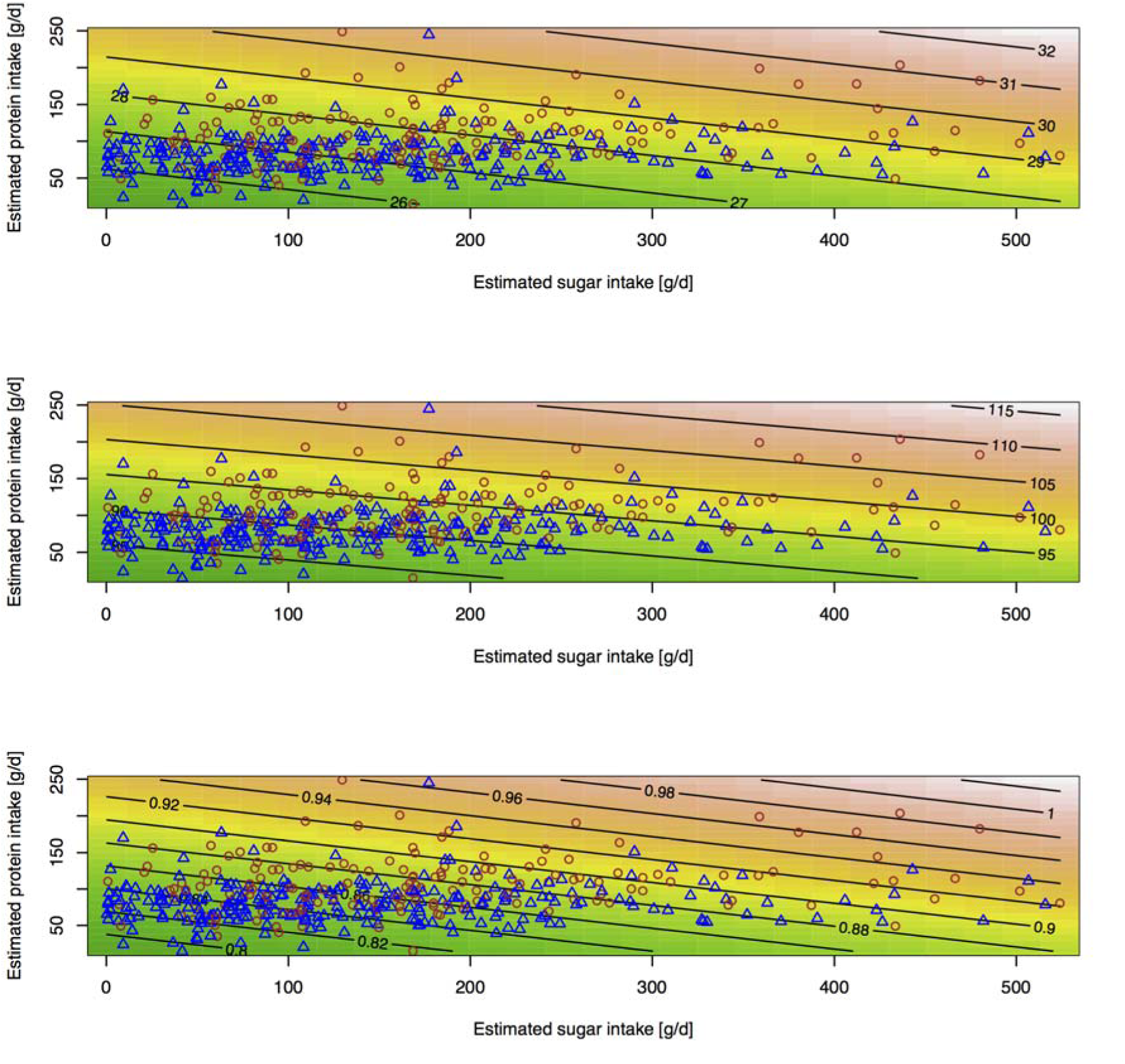
Association between estimated sugar, protein intake and obesity risk markers using a response surface model. Association between estimated total sugars and protein intake and (a) BMI [kg/m^2^], (b) waist circumference [cm] and (c) waist-to-hip ratio in women (blue triangles) and men (brown circles) using a response surface model. Points show data for individual participants, contour lines and colours estimated BMI, waist circumference and waist-to-hip ratio of linear regression mode respectively.

We have explored these relationships further by using uncalibrated biomarker data. Our data show a strong association between urinary sucrose and measures of obesity, as well as obesity risk based on waist circumference and waist-to-hip ratio. These associations were generally strengthened when including sucrose and fructose in the same model. Conversely, there were no significant associations for urinary fructose and only few associations were significant for total urinary sugars.

These results suggest that the associations between sugar intake and measures of obesity are mainly driven by sucrose. In contrast to fructose, which is derived from dietary fructose and hydrolysed sucrose and extensively metabolised, the only source of urinary sucrose is dietary sucrose^14-16,33^, making it more sensitive to changes in sucrose intake, the main contributor to intake of free sugars in the UK. Furthermore, high-fructose corn syrup (HFCS) or isoglucose was not commonly used in England at the time of the study as import and production was tighly controlled as part of the European Union sugar regime (Commission Regulation (EC) No 314/2002). Therefore the main source of dietary fructose were fruit and fruit products, such that fructose was most likely a surrogate marker of their intake.

Our results show that urinary sugars can be used to estimate sugar intake in the general population when 24h urine samples are available. In the context of current discussions regarding sugar intake and the recently updated WHO recommendations on sugars intake (1), the biomarker could be used to monitor the efficacy of public health interventions. Furthermore, we showed significant associations between sugar intake and BMI, confirming results of previous observations in EPIC Norfolk^4,19^. It is the first time that such an association has been shown in a nationally-representative sample of the general population using a validated biomarker. Our data also show significant associations between protein intake and measures of obesity and risk, however, in contrast to protein, sucrose is not an essential part of the human diet and intake can be reduced without adverse effects.

## 5 Acknowledgements

The technical assistance of K L Jones and D A Jones (IBERS Analytical Chemistry) is acknowledged for the nitrogen analysis.

